# Influence of native endophytic bacteria on the growth and bacterial crown rot tolerance of papaya (*Carica papaya*)

**DOI:** 10.1101/2019.12.22.886341

**Authors:** Mark Paul Selda Rivarez, Elizabeth P. Parac, Shajara Fatima M. Dimasingkil, Eka Mirnia, Pablito M. Magdalita

**Affiliations:** Institute of Plant Breeding, University of the Philippines Los Baños, Los Baños 4031, Laguna, Philippines; Department of Biotechnology and Systems Biology, National Institute of Biology, 1000 Ljubljana, Slovenia; College of Agriculture and Agri-Industries, Caraga State University, 8600 Butuan City, Philippines; College of Agriculture, Mindanao State University-Maguindanao Campus, 9601 Maguindanao, Philippines; Assessment Institute of Agricultural Technology-West Sumatera, Indonesian Agency of Agricultural Research and Development, 25001 West Sumatra, Indonesia

**Keywords:** papaya bacterial crown rot, *Erwinia*, *Kosakonia*, *Sphingomonas*, *Bacillus*, endophytic bacteria, autoclaved culture metabolites

## Abstract

The native plant microbiome is composed of diverse communities that influence its overall health, with some species known to promote plant growth and pathogen resistance. Here, we show the antimicrobial and growth promoting activities of autoclaved culture metabolites (ACMs) from native endophytic bacteria (NEB) in a papaya cultivar that is tolerant to bacterial crown rot (BCR) caused by *Erwinia mallotivora*. Initially, bacterial colonization in recovering tissues of this cultivar was observed before onset of tissue regeneration or ‘regrowth’. We further isolated and characterized these bacteria and were able to identify two culturable stem NEB under genera *Kosakonia* (EBW), related to Enterobacter, and *Sphingomonas* (EBY). We also identified root NEB (BN, BS and BT) under genus *Bacillus*. Inhibition assays indicated that ACMs from these NEB promptly (18-30h) and efficiently inhibited (60-65%) *E. mallotivora* proliferation *in vitro*. Interestingly, when ACMs from BN and EBW were inoculated in surface-sterilized papaya seeds, germination was variably retarded (20-60% reduction) depending on plant genotype, but plant biomass accumulation was significantly stimulated, at around two-fold increase. Moreover, greenhouse experiments show that ACMs from all isolates, especially EBW, significantly reduced BCR incidence and severity in susceptible genotype, at around two-fold. In general, our observations of pathogen antagonism, plant growth promotion leading to disease reduction by ACMs of native endophytic bacteria suggested its contribution to increased fitness of papaya and tolerance against the (re)emerging BCR disease.

## Introduction

Papaya (*Carica papaya*) is an economically important tropical fruit, mainly cultivated for food and cosmetic products. Due to its popularity, production of papaya steadily increased by 4.35% globally from 2002 to 2010 (Evans and Ballen, 2012). However, the industry still faces yield losses due to diseases, causing problems from plantation, transportation and processing until marketing (Ventura et al., 2004). Most importantly, emerging diseases greatly impair papaya production, and even cause up to 100% yield loss in some afflicted areas, especially in Asian tropical countries (Ploetz, 2004). A re-emerging disease of papaya, named bacterial crown rot (syn. soft rot, canker, decline) caused by a bacterium of genus *Erwinia* was reported in Southeast Asia (Maktar et al., 2008). Historically, the disease was reported in Java (von Rant, 1931), and in other regions such as Mariana Islands (Trujillo and Schroth, 1982) and in the US Virgin Islands (Webb, 1984). In Mindoro, Philippines, a similar soft rot disease, which has water-soaked symptoms on leaves, petioles and whorl accompanied by a foul odor was reported. The pathogen was identified under the genus *Erwinia* (Pordesimo and Quimio, 1965) and remained a major problem until the 1980s (Obrero, 1980).

Re-emergence of *Erwinia* rot in papaya was reported in many regions across the globe in recent past decade. In early 2000, the bacterial ‘canker’ disease of papaya was reported in the Mediterranean region, caused by *Erwinia papayae* sp. nov. (Gardan et al., 2004). In Asia, a bacterium identified as *Erwinia papayae* was reported to cause ‘papaya dieback’ disease in Malaysia (Maktar et al., 2008). The ‘crown rot’ disease that was recently observed in the Philippines (dela Cueva et al., 2017) and ‘black rot’ disease in Japan (Hanagasaki et al., 2020) were both reported to be caused by *Erwinia mallotivora*. Interestingly, the ‘crown rot’ disease of papaya in the Republic of Tonga in Africa was reported to be caused also by *E. mallotivora* (Fullerton et al., 2011). In the Philippines, studies were reported regarding the possible resistance or tolerance of papaya against the bacterial crown rot (BCR) disease. A slow hypersensitive response exhibited by tolerant genotypes were reported. This was followed by tissue hardening and lateral stem regrowth at the junction of the diseased and healthy stem in some genotypes. The phenomenon, referred to as ‘regrowth’, is thought to be the manner by which papaya recovers from the disease, and perhaps a mechanism for tolerance (Magdalita et al., 2016). Breeding for resistance and development of management strategies against the bacterial crown rot disease are already in its early stages in the Philippines. Regrowth hybrid lines with good marketable qualities and with resistance to papaya ringspot disease are currently under preliminary field trials and back-cross F3 progenies are now in the pipeline (Magdalita et al., 2015).

However, these candidate BCR-tolerant cultivars are still undergoing field trials and there is a great need to develop augmentative solutions to combat BCR emergence and spread. Fortunately, in recent years, research in the role of endophytic bacterial community in plant defense has improved greatly (Podolich et al., 2015). In a recent study, autoclaved culture metabolites (ACMs) from *Bacillus* species were shown to significantly induce disease resistance against *Pseudomonas syringae* pv. lachrymans in cucumber and pepper (Song et al., 2017). Moreover, growing evidences have shown the important role of plant endospheric microbiome in plant defense and overall fitness (Liu et al., 2020; Trivedi et al., 2020). Here, we showed the possibility of exploiting native endophytic bacteria (NEB) of papaya stem and roots, a possibility that was recently reported in NEBs from papaya seeds (Mohd Taha et al., 2019). We prepared ACMs as previously shown (Song et al., 2017) and investigated its effect on fitness of a susceptible cultivar, contributing to BCR tolerance. Specifically, we also performed antimicrobial assays using ACMs from NEBs and investigated in effect on seed germination, plant growth and response to *E. mallotivora* challenge. We hypothesize that antimicrobial activity, growth promotion and resistance-induction of ACMs from NEBs could be acquired or external mechanisms for pathogen defense in plants based. The use of these augmentative defense strategies in as early as seed or seedling stage were also suggested to help in developing more resilient genotypes or cultivars (Singh et al., 2018) that are otherwise highly susceptible to disease, but with good marketable qualities.

## Materials and Methods

### Ultrastructural observation and isolation of endophytic bacteria in regrowth tissues

Tissue samples from two-month old tolerant cultivar 5648 that were freshly recovering from BCR infection were prepared and examined using scanning electron microscope (SEM). Imaging was done using the Phenom XL™ (Phenom-World BV, Eindhoven, The Netherlands), a table top SEM. To further isolate observed bacteria, stem portions from the regrowth plants were directly obtained from the junction of the regrowth and the healthy stem part. Similarly, portions of the roots of regrowth plants tissues were prepared. The tissues underwent surface sterilization with following procedure: 95% ethanol for 1 min, 3% sodium hypochlorite solution for 6 min and 95% ethanol for 0.5 min followed by rinsing twice with sterile distilled water for 5 min (Coombs and Franco, 2003). Thereafter, sterilized tissues were submerged for 10 min in 5 mL sterile water in a test tube to visibly detect bacterial ooze. A loopful of the serially diluted (10^−8^) bacterial suspension was streaked onto peptone sucrose agar (PSA) plates and incubated at room temperature for 24 h. Single colonies of actively growing bacteria were transferred to new PSA plates for subculture and purification. Pure colonies of the bacteria were maintained in PSA culture slants or as glycerol-skim milk stocks and stored at 4°C for future use.

### Cultural and biochemical characterization and pathogenicity tests of bacterial isolates

All cultural characterization tests conducted in this part of the study were adapted from the procedures in phytobacteriology books (Goszczynska et al., 2000; Borkar, 2017) with minor modifications. In order to prepare single colonies, a loopful of a serially diluted suspension (10^−8^) of the isolates was streaked multiple times onto the surface of PSA plates. The color, colony diameter, morphology (*e*.*g*. shape, margin and elevation) and consistency were noted. Additionally, single stroke colonies were prepared in nutrient agar (NA) and described after one week. Colony color, shape, and other characteristics were likewise noted. Additionally, growth in nutrient broth was characterized in a test tube, where observations were done after one and two weeks. The morphology of the bacterial cells was characterized using gram staining methods. Biochemical tests include assay for catalase, cysteine desulfurase, lipase and gelatinase activities, while physiological test include test for oxygen requirement and utilization of starch and other sugars. All tests conducted in this part of the study were adapted from the procedures in phytobacteriology books (Goszczynska et al., 2000; Borkar, 2017) with minor modifications. Identification based on biochemical tests were done using the *Bergey’s* Manual of Systematic Bacteriology (Whitman et al., 2015). The tests were done in triplicates and the entire experiment was repeated twice to verify results.

On the other hand, in order to test for pathogenicity of the isolates, suspensions of the bacteria grown PSA slants were used in inoculation of two-month old seedlings at a concentration of 10^8^ cfu/mL. BCR-susceptible genotype 336 was used as the test plant. The plants were wounded first at the leaf petiole and leaf midrib using a sterile needle. The inoculum was delivered using a sprayer at approximately 1.0 mL per plant. Observations were done at 7 and 21 d after inoculation. Successful infection is indicated by necrosis and water-soaked lesions and progressive rotting from the point of wounding.

### DNA extraction and amplification of 16S rRNA

DNA extraction was done using a simple protocol developed by Chen and Kuo, with minor modifications (Chen and Kuo, 1993). Single pure colonies were harvested from 72-h slant cultures in PSA. Both genomic DNA and diluted colony cell suspension were used as template for PCR reactions to amplify 16S rRNA of the bacterial isolates. Around 10-20 ng of DNA was used as template in PCR reactions and around 10-20 cells/mL were used as template in cell suspension PCR. For the 25 uL PCR reaction mixture, the final concentration of PCR reagents (Invitrogen, MA, USA) was as follows: 1X PCR buffer without MgCl_2_, 2.0 mM MgCl_2_, 0.2 mM dNTPs, 0.5 μM of forward primer and 0.5 μM reverse primer. *Taq* DNA polymerase was added at 0.04 U/μL. Universal 16S rRNA primers used were F8/rP2 and 27F/1492R (Weisburg et al., 1991). The PCR program was set at 94°C for 10 min for initial denaturation, followed by 32 cycles of 94°C for 30 s, 55°C for 40 s and 72°C for 60 s, and final extension 72°C for 5 min and indefinitely at 4°C for storage. PCR amplicons were checked for correct size, at around 1300 bp, and purity in 1% agarose gel. Amplicons were then sent for Sanger sequencing at AIT Biotech, Inc. (Singapore).

### Bioinformatic analysis using 16S rRNA to identify bacterial isolates

The bacterial isolates were identified up to the closest taxa using 16S rRNA sequence. For the purpose of identifying closest species-level taxon, we primarily used valid species (*i*.*e*. type specimens) curated in EZ-BioCloud (www.ezbiocloud.net/identify), a public data and analytics for the taxonomy of bacteria and archaea (Yoon et al., 2017). The sequences of closest accessions were collected and used further in phylogenetic analysis in Molecular Evolutionary Genetics Analysis software version 7 (MEGA7) (Kumar et al., 2016). Multiple sequence alignment following the ClustalW method (Thompson et al., 1994) was performed before inferring phylogenetic tree using Neighbor-Joining method (Saitou and Nei, 1987) following the Tamura-Nei model (Tamura and Nei, 1993). Lastly, we also used BLAST (Basic Local Alignment Search Tool) in NCBI (https://blast.ncbi.nlm.nih.gov/Blast.cgi) (Altschul et al., 1990) to check for the identity of our isolates to closest species or strains in GenBank.

### Antimicrobial assay using autoclaved culture metabolites (ACMs)

A suspension of the 72-h old culture (log phase) of the endophytic bacteria and *Bacillus* isolates in PSA was diluted to obtain approximately 10^8^ cfu/mL or OD_600_ at 0.75-0.80. A 1.0 mL aliquot consisting of approximately 10^8^ cells was inoculated in a 100 mL 3% tryptic soy broth (TSB, BD-Difco) in Erlenmeyer flasks. The broth cultures were incubated for 72 h at room temperature with continuous shaking at 120 rpm. Thereafter, the bacterial broth cultures were incubated for 30 min at autoclaving temperature (121°C) and pressure (15 psi) in order to retain only the autoclaved culture metabolites (ACMs). The ACMs were centrifuged at 12,000 rpm for 10 min to separate cell debris. The supernatant was then filter-sterilized using a 0.2 µm-pore size polyether sulfone (PES) membrane filter (Sterile Acrodisc, Gelman Sciences or Whatman, GE Life Sciences) to remove remaining bacterial cell debris. To use ACM filtrates for *E. mallotivora* inhibition assays, dilutions were made to obtain approximately OD_600_ of 0.25 to 0.30 as standard concentrations.

Log phase colonies (72-h old) of *E. mallotivora* was prepared as suspension in sterile water at 0.25 to 0.30 OD at 600 nm, in order to obtain approximately 10^4^ cells/mL. The assay was done in a 96-well microtiter flat-bottom polycarbonate plate (Tarsons, Cat. No. 941146). ACMs and *E. mallotivora* cells were mixed in a well at 1:1 ratio (50 µL + 50 µL). Wells that served as controls contain *E. mallotivora* cells only and *E. mallotivora* cells in tryptic soy broth while tryptic soy broth and sterile water only served as blanks. The OD_600_ reading and log_10_ cfu/mL of the wells at 0, 24, 48 and 72 h were noted and compared across treatments. In order to have a good estimate of the inhibitory effect of the ACMs, percent inhibition was calculated using the formula: 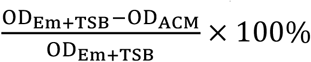, where OD_Em+TSB_ is the optical density reading of the negative check (Em+TSB) at a given time point and OD_ACM_ is the optical density of any ACM-treated *Em* suspension at a given time point. This assay followed a randomized complete block design (RCBD), wherein the ACMs served as treatments and replicates as the blocking factor. Three replicates were prepared for each treatment containing three samples with 12 subsamples. Thus, for each treatment, 96 readings were done every time the OD values were assessed.

### Preparation of seeds and ACM treatment

The papaya seeds used in this study were obtained from the papaya germplasm collection at the Fruit Breeding Laboratory at Institute of Plant Breeding, College of Agriculture and Food Science, University of the Philippines Los Baños. The genotypes used were variety *Cariflora* (genotype 5648), a PRSV-tolerant and BCR-tolerant ‘regrowth’ genotype with good yield and fruit quality, and the *Solo* variety (genotype 501), which is susceptible to both PRSV and BCR. Sterilized coir dust-garden soil medium was prepared as potting medium for the seeds. Seeds were surface sterilized in an Erlenmeyer flask using 70% ethanol solution for 1 min, followed by rinsing twice in sterile water for 2 min and blot- and air-drying under the laminar flow hood for 15 min. The seeds were immersed in the ACMs for 16 h, followed by blot- and air-drying under the laminar flow hood for 30 min. Soaking in sterile tryptic soy broth only served as the mock treatment. The seeds were sowed in sterile potting medium with complete (40-40-40) fertilizer and maintained under greenhouse conditions with daily watering for six weeks. The plants were ready for inoculation if it has 4-5 leaf nodes already and about 10 cm in height. The effects of ACM treatment on seed germination was assessed based on percent of seeds germinated after four weeks. To evaluate the effect of ACM treatment on plant biomass, the 7-wk old ACM-treated plants were slowly dried in an oven at 65°C. The dry weight of the samples were assessed after 36 h. This experiment followed a randomized complete block design (RCBD), wherein the ACMs served as treatments and replicates as the blocking factor. Separate statistical analyses were done for the two genotypes.

### Inoculation of ACM-treated plants and disease assessment

Six-week old ACM-treated seedlings of both papaya genotype 5648 and 501 were inoculated with *Erwinia mallotivora* as described by Supian *et al*., with some modifications (Supian et al., 2017). The plants were watered first before inoculation to increase its internal turgor pressure and activate transpiration and water translocation, thus increasing inoculation efficiency. To further increase chances of infection, the apical tips, petiole and midrib of the second and third youngest leaf were pricked with a sterile syringe needle. On the other hand, the inoculum was prepared by suspending a 72-h old pure culture of *E. mallotivora* in sterile water and diluting the suspension to obtain approximately 10^8^ cfu/mL (OD_600_ of 0.75-0.80). The inoculum was applied through spraying three times per plant, delivering approximately 1.0 mL of inoculum. After inoculation, plants were covered with clear plastic cups in order to maintain high relative humidity and favor bacterial growth for 48 h.

Disease assessment was done at 7 and 21 days post inoculation (dpi). Percent disease incidence was calculated as 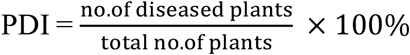. Disease severity assessment followed the scoring used by Supian *et al*. (2017) with some modifications. This modified scale consisted of the following: 0 (no symptom), 1 (lesions are restricted to inoculation point only), 3 (lesions are expanding to less than half of leaf area), 5 (lesions are expanding to more than half of leaf area), 7 (lesions and rotting spread to the petiole and stem and 9 = (plant suffers severe rotting, crown falls off). Finally, percent disease severity index was calculated as (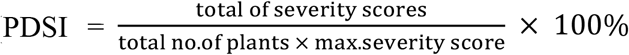. This assay followed a randomized complete block design (RCBD), wherein the ACMs served as treatments and replicates as the blocking factor. Three replicates were prepared for each treatment containing three samples with five subsamples each. Thus, for each treatment, 45 plants were allotted.

### Statistical analysis of data

All *in vitro* experiments were done twice and greenhouse experiment were done thrice to confirm results. All data generated were subjected to statistical analysis through one-way analysis of variance (ANOVA at α= 0.05) in SAS software program version 14.3 (SAS Institute Inc., Cary, NC). Separate statistical analyses were done for the two papaya genotype used. The means were subjected to pairwise mean comparison using Tukey’s honest significant difference (HSD) test. All tests are considered significant if *P* < 0.05 (*), *P* < 0.01 (**) or *P* < 0.001 (***).

## Results

### Native endophytic bacteria from ‘regrowth’ BCR-tolerant plants

Initially, a community of bacterial cells was observed in the tolerant genotype, 5746 (var. *Cariflora*) in early stages of regrowth (3-4 wks), but not in fully regrowth tissues (7-8 wks) (see Supplementary Figure 1). Building on the SEM observations, regrowth-associated culturable native endophytic bacteria (NEB) were isolated. Two morpho-culturally distinct, stem NEBs (EBW, EBY) and three root NEBs (BN, BS, BT) were isolated. Based on *Bergey’s* Manual of Systematic Bacteriology (Whitman et al., 2015), EBW was identified under genus *Enterobacter* (*Gammaproteobacteria*: *Enterobacterales*: *Enterobacteriaceae*). EBY was identified under genus *Sphingomonas* (*Alphaproteobacteria*: *Sphingomonadales*: *Sphingomonadaceae*). On the other hand, the *Bacillus* isolates were not easily identified using phenotypic characteristics because their colonies are very much alike and also exhibit similar biochemical properties. Identification using 16S rRNA sequence was carried out to determine the most probable taxa of the bacterial isolates. Figure 2 shows the resulting phylogenetic trees with the percent identity to valid taxa or type specimen identified in EzBioCloud. The analyses showed that EBW is paraphyletically related to *Kosakonia* sp. clade and monophyletically related the *Enterobacter* sp. clade. It is worthwhile to note that *Kosakonia* sp. is a new genus of plant*-*associated, endophytic ex. *Enterobacter* species (Brady et al., 2013). Moreover, EBY is paraphyletically related to *S. endophytica* YIM 65583 (Huang et al., 2012) and *Sphingomonas phyllosphaerae* FA2 (Rivas et al., 2004), although percent identity to closest taxa is not convincingly high enough, which also implies that it could be a new species (see Supplementary Table 1). On the other hand, the *Bacillus* isolates cannot be definitively identified because of inherent limitation of 16S rRNA for *Bacillus* taxonomy (Maughan and Van der Auwera, 2011). Nevertheless, it is possible that the three isolates are members of the *Bacillus subtilis* species complex (Fan et al., 2017) based on bioinformatic and phylogenetic analysis done. Lastly, all the isolates were not able to infect papaya nor cause any hypersensitive reaction, which implies no interaction with plant resistance machinery. The summary of characteristics of the isolates is shown in Table 1, while all supplementary data can be accessed in the through provided herewith.

**Table 1.** Summary of morpho-cultural and biochemical properties of the bacterial isolates isolated in this study.

| Properties | *E. mallotivora* | *Kosakonia* sp. | *Sphingomonas* sp. | *Bacillus* sp. BN | *Bacillus* sp. BS | *Bacillus* sp. BT |
| --- | --- | --- | --- | --- | --- | --- |
| **Cell / Single Colony<sup>1</sup>** |  |  |  |  |  |  |
| colony color | cream white | white | deep yellow | light brown | cream white | cream white |
| cell shape | rods | rods | short rods | rod | rod | rod |
| gram stain | negative | positive | negative | positive | positive | positive |
| spore formation | absent | present | absent | present | present | present |
| diameter (mm) | 0.8-1.0 | 6.0-6.5 | 5.5-6.0 | 6.0-6.5 | 7.0-7.5 | 5.0-6.0 |
| shape | circular | circular | circular | irregular | irregular | irregular |
| margin | entire | entire | entire | undulate | undulate | undulate |
| elevation | convex | convex | convex | umbonate | flat | raised |
| consistency | fluidal | fluidal | butterous | dry | dry | dry |
| **Morpho-cultural<sup>2</sup>** |  |  |  |  |  |  |
| nutrient agar stroke | filiform, butyrous | filiform, viscid | filiform, butyrous | effuse, butyrous | effuse, rubbery | rhizoid, brittle |
| nutrient broth | flocculent | ring | flocculent | membranous | membranous | membranous |
| **Biochemical<sup>3</sup>** |  |  |  |  |  |  |
| oxygen requirement | facultative anaerobe | facultative anaerobe | facultative anaerobe | obligate aerobe | obligate aerobe | obligate aerobe |
| catalase | + | + | + | + | + | + |
| cysteine desulfurase | - | + | + | + | - | - |
| lipase | - | + | + | + | + | + |
| gelatinase | - | + | - | + | + | + |
| starch utilization | + | - | +/- | + | + | - |
| cellobiose utilization | - | + | + | + | + | + |
| glucose utilization | + | + | + | + | + | + |
| lactose utilization | - | + | + | + | + | + |
| maltose utilization | - | + | + | + | + | + |
| mannitol utilization | + | + | - | + | + | + |
| mannose utilization | + | + | + | - | - | - |
| sorbitol utilization | - | + | - | + | + | + |

<sup>1</sup>Colony morphology in Peptone Sucrose Agar (PSA) after 7 d of room temperature incubation.
<sup>2</sup>Morpho-cultural properties after 14 d of room temperature incubation.
<sup>3</sup>+, positive test; -, negative test; +/-, weak positive.

**Figure 2.**
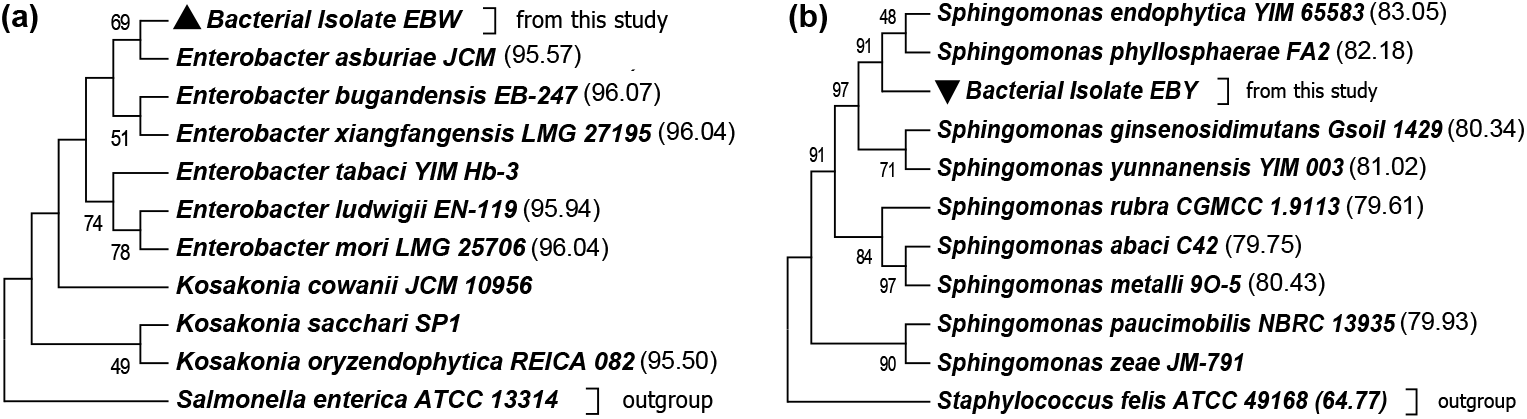
Phylogenetic trees showing most probable taxa and species-level identity based on 16S rRNA sequences of: (a) Isolate EBW tree showing relationship with *Enterobacter/Kosakonia* clade, percent identities to EBW are shown in the members of the clade; (b) Isolate EBY tree showing relationship to the *Sphingomonas* species, percent identities to EBY are shown in the members of the clade. The analysis followed Neighbor Joining following the Tamura-Nei model in MEGA7. Bootstrap values for 1000 replicates greater than 45% are shown in the nodes.

### *E. mallotivora* was promptly and efficiently inhibited by ACMs

The microtiter plate assay for the inhibition of *Em* was proven to be useful in evaluating inhibition efficacy of the autoclaved culture metabolites (ACMs) by just monitoring the absorbance readings. Figure 3 shows the trend in absorbance of the ACM treatments, as well as percent relative inhibition. After 24 h, a drastic increase in cell density was observed in the BS, BT and EBY treatments. Nevertheless, this observed increase in cell density in the ACM treatments was still considerably lower than the negative control, at around 2-3 folds. Thus, there is still a significant inhibition of cell multiplication in these treatments. More importantly, bacterial growth was highly inhibited when *Em* was subjected to BN and EBW at 60-65% and 53%, respectively.

**Figure 3.**
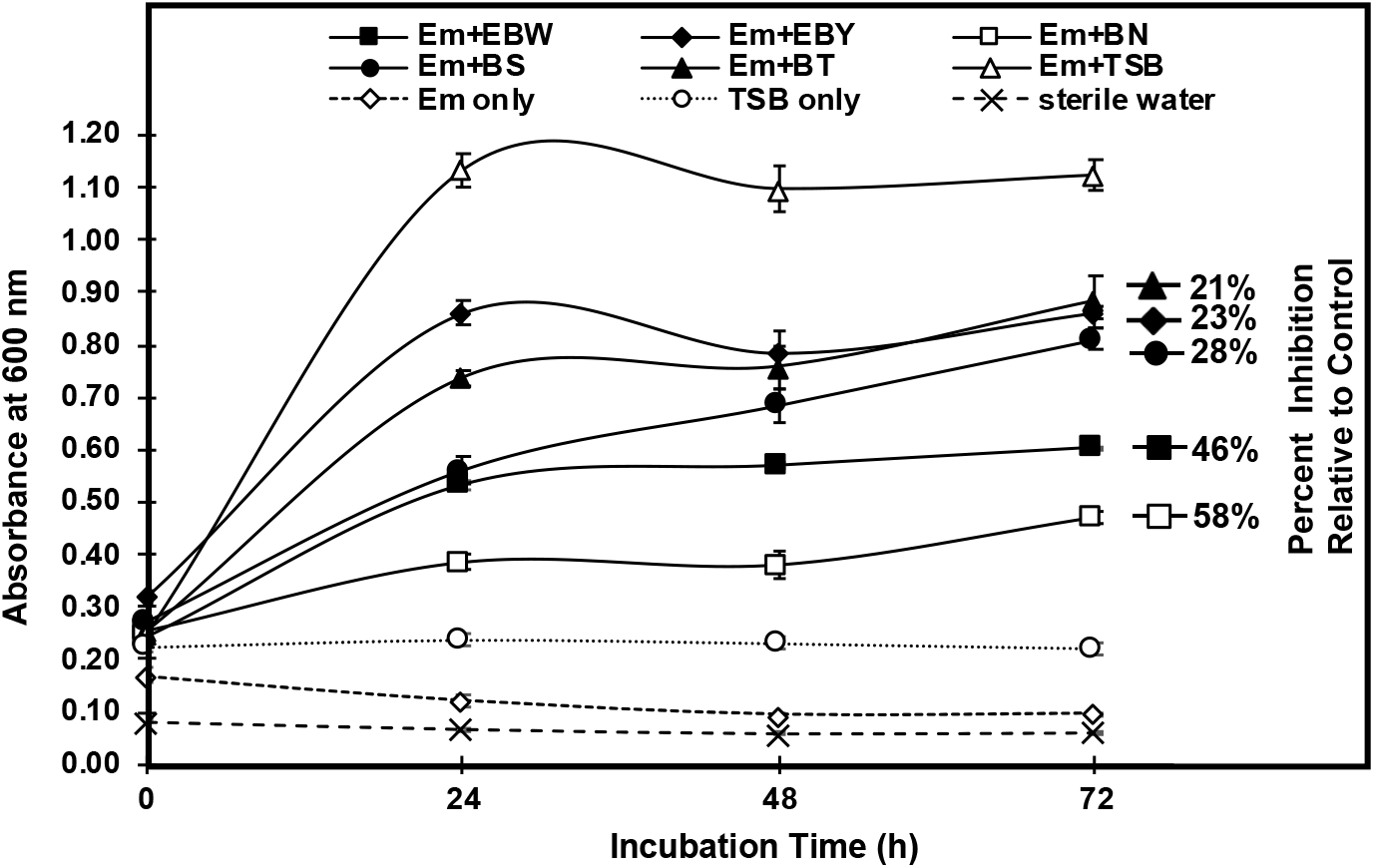
Antimicrobial effect of autoclaved culture metabolites (ACMs) on *Erwinia mallotivora* proliferation *in vitro*. Changes in absorbance readings at 600nm (left axis) through time (horizontal axis) and percent inhibition relative to the controls (right axis) of the autoclaved culture metabolite-treated *Erwinia mallotivora* (Em) cell suspensions are shown. The treatments are as follows: Em treated with ACM from isolate EBW (*Kosakonia* sp.) (Em+EBW), EBY (*Sphingomonas* sp.) (Em+EBY), isolate BN (Em+BN), isolate BS (Em+BS) and isolate BT (Em+BT). Em in tryptic soy broth (Em+TSB) and Em only served as negative controls while TSB only and sterile water only served as blanks. Standard deviation shown as error bars are depicted in each data point.

### All ACMs reduced growth vigor of papaya, except ACM from *Kosakonia*

Treatment of surface-sterilized papaya seeds with ACMs from bacterial endophytes and *Bacillus* isolates resulted to retarded germination compared to mock-treated seeds. Figure 4a shows the average percent germination of papaya seeds of genotype 501 (var. *Solo*) and 5648 (var. *Cariflora*), which were evaluated as susceptible and tolerant (*i*.*e*. regrowth) to the crown rot disease, respectively. Among the all treatments, BS has no significant adverse effect on the germination of papaya seeds in both genotypes. However, treatment with isolate BN and BT resulted to a significant reduction in percent germination of 501, while treatment with isolate EBW and EBY resulted to lowest percent germination in genotype 5648. This implies that papaya might respond differently to ACM treatments, depending on genotype. Plant growth was assessed by getting the dry biomass of plants at 7 wks after sowing. Figure 4b shows the accumulated biomass, measured as oven-dried weight, of the ACM-treated plants. Treatment with ACM from EBW yielded the highest dry weight, particularly in genotype 5648, which correspond to more than two-fold increase relative to mock. However, no significant increase in biomass was observed in genotype 501 in the same treatment, indicating that growth stimulation by bacterial ACMs in papaya might be genotype-dependent. Finally, all other treatments significantly reduced biomass accumulation in both genotypes as depicted, especially in BS ACM-treatment.

**Figure 4.**
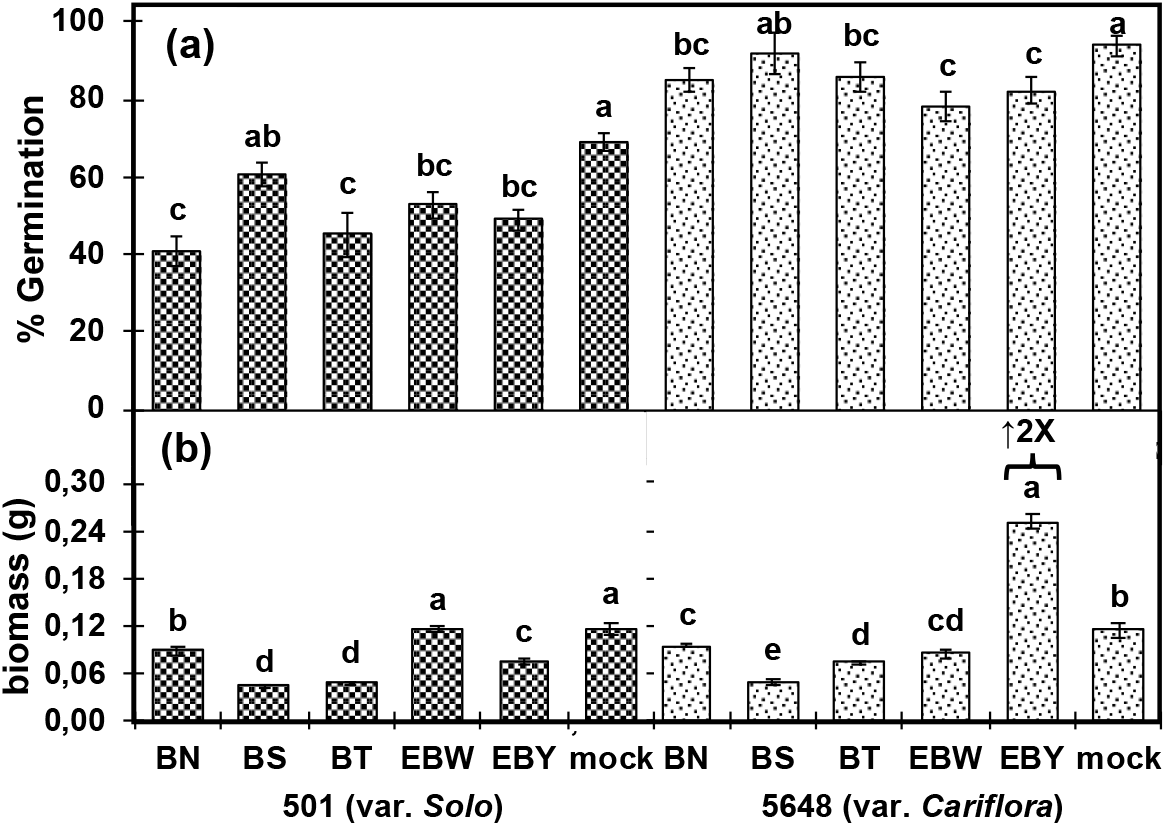
Effect of autoclaved culture metabolites (ACMs) on the growth vigor of papaya. (a) Germination data taken at 21 days after sowing (das) and (b) biomass taken at 49 das. Columns with checkered shade are data from the papaya BCR-susceptible genotype and columns with dotted shade are data from papaya BCR-tolerant genotype. The two-fold increase in dry biomass is EBY-treated genotype 5648 is also highlighted. Treatments marked with different letters are statistically different after HSD test at *P* < 0.05 (*).

### ACMs significantly reduced the incidence and severity of BCR disease in Papaya

The possibility of using easy to prepare, sterile, highly-stable, and potent metabolites from beneficial bacteria as promoter of disease resistance in papaya was assessed in this study. Although plant vigor was adversely affected by the treatment of ACMs, the papaya plant would eventually recover and grow normally. Figure 5a,b shows the representative photographs of inoculated plants after two weeks of observation. Based on observations made, BN-treated plants in both genotypes seemed to have retarded growth after 14 dpi, due to infection. Necrosis within half of the inoculated leaf area was commonly observed in these plants. BS- and BT-treated plants in both genotypes were not significantly affected by the pathogen challenge and continued to grow normally. At most instances, plants in this treatment only had necrosis at the inoculation point. Recorded percent disease incidence (Figure 5c) revealed that in general, ACM-treated plants have lower propensity to bacterial crown rot infection. BCR incidence continued to increase until 21 dpi, but the incidence in ACM-treated plants remained lower compared to that of the mock-treated plants, in both genotypes. On the other hand, there was a general reduction in average severity ratings in both genotypes (Figure 5d). More importantly, a two-fold reduction in disease severity was observed in EBW treatment at 21 dpi relative to control. Lowest severity was observed in EBW-treated 5648 plants relative to control. Generally, improved resistance is more pronounced in 5648 (resistant) plants than in 501 (susceptible) plants.

**Figure 5.**
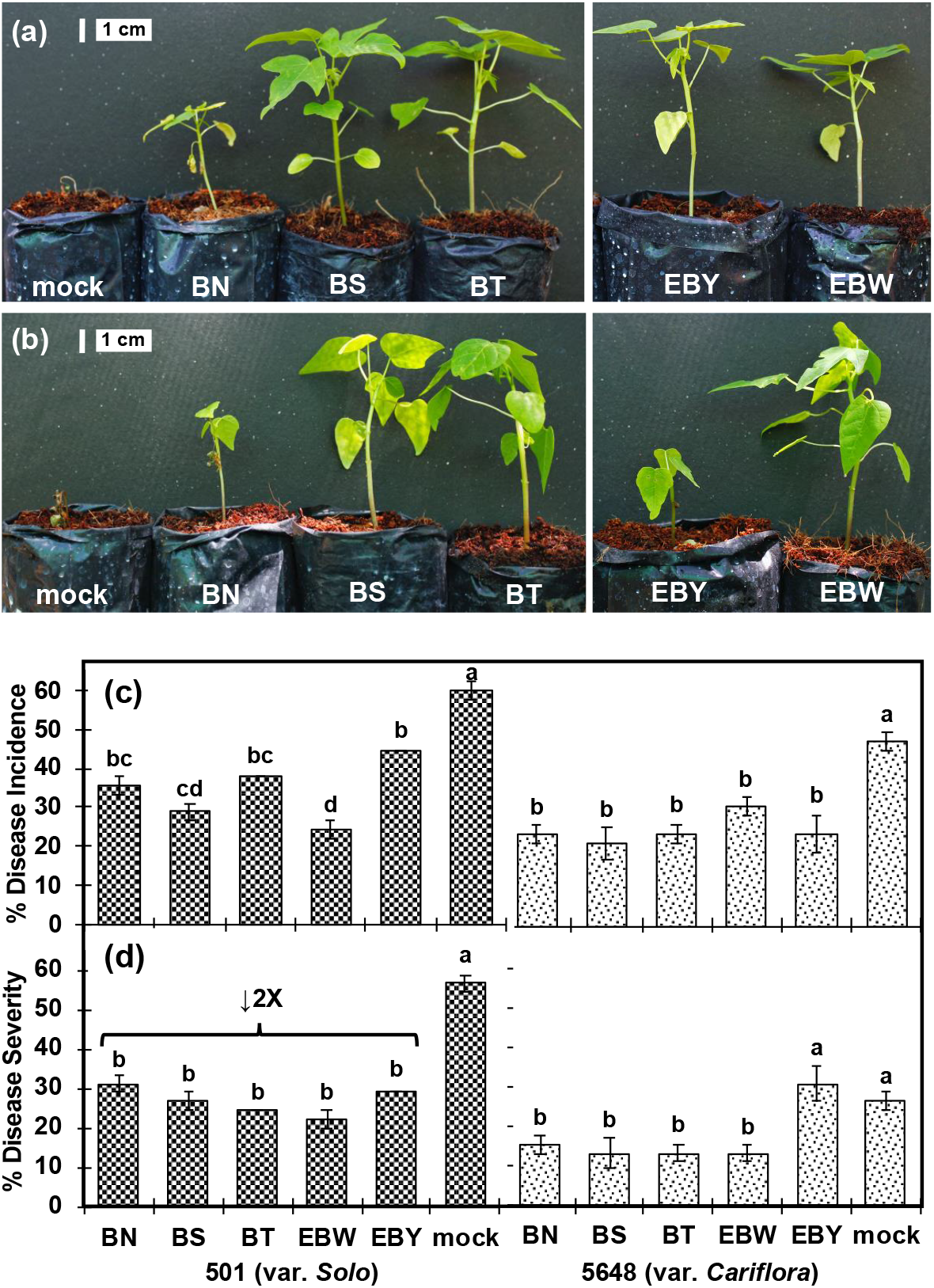
Effect of autoclaved culture metabolites (ACMs) on the incidence and severity of bacterial crown rot disease of papaya. Representative photos of pathogen-challenged ACM-treated plants of (a) genotype 501 and (b) genotype 5648. Scale bars are shown in the figure above. (c) Average percent disease incidence taken at 21 days after inoculation (dai), (d) average percent disease severity at 21 dai. Columns with checkered shade are data from the papaya BCR-susceptible genotype and columns with dotted shade are data from papaya BCR-tolerant genotype. Treatments marked with different letters are statistically different after HSD test at *P* < 0.05 (*).

## Discussion

BCR tolerant genotypes such as 5648 (var. *Cariflora*) were primarily identified using the lateral stem ‘regrowth’ phenotype at 5-7 wks after infection (wai) as indicator of disease resistance or tolerance. In this study, it was observed through SEM that infection sites in recovering tissues from genotype 5648 were colonized by a community of bacteria at the onset of regrowth (3-4 wai). Building around the initial observations, endophytic bacteria from stem and roots were isolated from regrowth genotype 5648, and further characterization and identification using 16S rRNA were carried out thereafter. Root-associated *Bacillus* species were identified under the *Bacillus subtilis* species complex, while the two endophytic bacteria were identified under genus *Kosakonia* ex. *Enterobacter* (isolate EBW) and *Sphingomonas* (isolate EBY). Related species under the *B. subtilis* species complex include *Bacillus velezensis* strain CR-502 (isolate BN) that was shown to produce surfactants that confer antimicrobial activity against other microbes (Ruiz-García et al., 2005). Additionally, antibiotics from *Bacillus* successfully inhibited gram-positive bacteria that are also human-pathogenic such as *Stapylococcus aureus, Clostridium difficile* and *Listeria monocytogenes* (Collins et al., 2016). We report herewith the first instance that a strain of the *B. subtilis* species complex was shown to have inhibitory properties against a bacterial plant pathogen.

Species of the genus *Sphingomonas* are reported to have beneficial properties and are usually associated as endophytes in many plants. In particular, *S. paucimobilis* associated with an orchid plant (*Dendrobium officinale*), was proven to have growth promoting properties by increasing phytohormone production and aiding in atmospheric nitrogen fixation, processes which the plant cannot carry out normally (Yang et al., 2014). Likewise, *S. phyllosphaerae* and *S. endophytica* were originally isolated as endophytes in *Acacia caven* and *Artemesia annua*, respectively. On the other hand, *Kosakonia* sp. (formerly *Enterobacter* sp.) is a new taxon name assigned to endophytic, plant-associated *Enterobacteraceae* species. Two species, *K. oryzophilus* and *K. oryzendophytica*, were found to be key inhabitants of rice root endosphere (Hardoim et al., 2013). The two species were able to promote rice growth by supplying nitrogen and phosphorus to the plant. Similarly, *Kosakonia sacchari* (Zhu et al., 2013) (formerly *Enterobacter* sp.) is an endophyte associated with sugarcane stem, roots and rhizosphere. The bacterium was shown to help the plant in producing fixed nitrogen for its consumption. Another species of *Kosakonia*, named *K. pseudosacchari*, was discovered to be endophytically associated with field grown corn (*Zea mays*) root tissues. *K. pseudosacchari* is also a closely related species of *K. sachari* and *K. oryzedophytica* (Kämpfer et al., 2016). However, there were no studies were reported yet regarding the antimicrobial properties of *Kosakonia* and *Sphingomonas* species other than this study. Interestingly, the aforementioned *Bacillus* species and endophytic bacteria were not able to infect papaya plants nor cause any hypersensitive reaction and is resistant to latex exposure. A similar set of endophytic bacteria were also isolated from and constantly associated with tissue-cultured papaya plants based on previous studies (Thomas et al., 2007b, 2007a; Thomas and Kumari, 2010).

Endophytic bacteria and various *Bacillus* species associated with plants were reported to have growth-promoting and disease resistance-inducing properties (Hallmann et al., 1997; Ongena and Jacques, 2008; Podolich et al., 2015). Species of *Bacillus*, such as *B. paralicheniformis* MDJK30 was reported to promote growth in *Arabidopsis thaliana* (Dunlap et al., 2015; Palacio-Rodríguez et al., 2017) and suppress growth of *Fusarium solani* (Wang et al., 2017). Another strain, *B. paralicheniformis* KMS 80 was discovered to be associated with rice root endosphere and helped the plant in nitrogen fixation (Annapurna et al., 2018) (Annapurna *et al*., 2018). However, assays for growth-promotion and disease resistance induction heavily rely on the availability of culturable isolates and its compatibility or ability to colonize the host plant. In this study, these challenges were surpassed by preparing culture metabolites that are easy-to-prepare, sterile and highly stable, thus the term, autoclaved culture metabolites (ACMs). Moreover, seed inoculation using ACMs of bacteria is a relatively new method used in stimulating plant growth and in turn, enhance disease resistance in plants. This method was previously demonstrated in pepper and cucumber using ACMs of *Bacillus gaemokensis* PB69 (Song et al., 2017). Unlike the study by Song *et al*., this study used unpurified preparations of ACMs, which contains mixture of metabolites that might have caused reduced germination in papaya seeds. However, it was shown that growth vigor of papaya seedlings was enhanced later on as shown in higher biomass accumulation compared to the normal or untreated ones. Comparatively, treatment of live cell suspensions of endophytic and other beneficial bacteria also led to reduced germination and seedling vigor in papaya (Thomas et al., 2007a), tomato (Worrall et al., 2012) and oil palm (Azri et al., 2018), but recovery of plant growth was observed when grown in greenhouse conditions. Nevertheless, live cell suspensions are prone to contamination and the cells are unprotected and unstable unlike when using ACMs.

On the other hand, ACMs from the NEBs contributed to the reduction of the incidence of crown rot, in both susceptible and resistant genotypes. The plants remained healthy wherein only local necrosis at the inoculation site was observed in the test plants. These observations indicated that ACM-treatment has a potential to decrease the predisposition of the papaya plant to the crown rot disease, as a collateral effect of plant growth stimulation. However, these observations still demand further molecular transcriptomic investigation to be referred to as a disease biopriming phenomenon. Interestingly, genotype dependence of papaya responses to ACM treatments might be important to consider in breeding programs to augment disease resistance. Endophytic bacteria and many *Bacillus* species were proved to have such bioactivity in plants, may it be its native host or a non-host (Tonelli et al., 2011; Yang et al., 2014; S. et al., 2018; Wicaksono et al., 2018). However, induction of defense responses in plants comes with a penalty or cost (Cipollini et al., 2003; Dietrich et al., 2005) by delaying and lowering the plant’s germination and seedling vigor. Nonetheless, we show that fully-grown seedlings eventually recuperate and even accumulate higher biomass relative to the untreated counterpart.

The use of live cell suspension remains the most popular method in exploiting beneficial microbes to promote better plant health (Hallmann et al., 1997). However, we present herewith, the use of autoclaved culture metabolites (ACMs) is relatively new but promising and only a few studies were reported to prove resistance-inducing bioactivity in plants (Noh et al., 2017; Song et al., 2017). The use of ACMs provides a convenient and fast way to screen plant-associated microbes because this method is easy to prepare, and the metabolites produced are highly stable and potent. If optimally applied, disease protection and induction of systemic resistance using beneficial microbes or its metabolites will be a good alternative to chemical pesticides or resistance inducers.

## Acknowledgements

The authors would like to thank Valeriana Justo, Lorele Trinidad, Ireneo Pangga, Teresita Dalisay and Fe dela Cueva of UPLB for their technical support in the conduct of the experiments.

## Supplementary Data

All supplementary data for this study can be downloaded through this link: https://doi.org/10.6084/m9.figshare.13499409.v1

## Declarations

### Availability of data and material

Sequences of the 16S rRNA gene for each bacterial isolate are deposited in NCBI GenBank. Accession numbers and sequences are also available in supplementary data [link].

### Author contributions

MPSR and PMM did the study conception and design and funding acquisition. PMM supervised all experiments. Material preparation and data collection were performed by MPSR, EPP, SFMD and EM. MPSR did the data analysis. The first draft of the manuscript was written by MPSR and all authors approved the final manuscript.

## Compliance with Ethical Standards

### Research involving Human Participants and/or Animals

We confirm that in this research any human and/or animals participant was not used and there is no any disagreement with informed consent.

## Supplementary material

202012_Supplemetary Figure 1 and Table 1.docx

ALL FINAL SEQUENCES.fa.txt

202012_Supplementary Data_EZ-BioCloud hits.xlsx

